# Tviblindi algorithm identifies branching developmental trajectories of human B cell development

**DOI:** 10.1101/2024.01.11.575178

**Authors:** Marina Bakardjieva, Jan Stuchlý, Ondřej Pelák, Marjolein Wentink, Hana Glier, David Novák, Jitka Stančíková, Daniela Kužílková, Iga Janowska, Marta Rizzi, Mirjam van der Burg, Tomáš Kalina

## Abstract

Detailed knowledge of the human B-cell development is crucial for proper interpretation of inborn errors of immunity and for malignant diseases. It is of interest to understand the kinetics of protein expression changes during the B cell development, but also to properly interpret the major and possibly alternative developmental trajectories. We have investigated human bone marrow and peripheral blood samples from healthy individuals with the aim to describe all B-cell developmental trajectories across the two tissues. We validated a 30-parameter mass cytometry panel and demonstrated the utility of “*vaevictis*” visualization of B-cell developmental stages. We used our recently developed trajectory inference tool “*tviblindi*” to exhaustively describe all trajectories leading to all developmental ends discovered in the data. Focusing on Natural Effector B cells, we demonstrated the dynamics of expression of nuclear factors (PAX-5, TdT, Ki-67, Bcl-2), cytokine and chemokine receptors (CD127, CXCR4, CXCR5) in relation to the canonical B-cell developmental stage markers (CD34, CD10, sIgM, IgD, CD20, CD27). Lastly, we performed analysis of the expression changes related to developmental branching points (Natural Effector versus Switched Memory B cells, marked by up-regulation of CD73).

In conclusion, we developed, validated and presented a comprehensive set of tools for investigation of B-cell development.

## Introduction

B-cells, together with T-cells, are adaptive immunity constituents responsible for antigen-specific responses and immune system memory. Mature and terminally differentiated B-cells produce high affinity antibodies. B-cells develop from hematopoietic stem cells in the bone marrow, exit to peripheral blood, enter the secondary lymphoid organs upon antigen encounter to mature in the germinal center and recirculate to peripheral blood and eventually home back to the bone marrow as antibody secreting cells.

These principles, key developmental stages and molecular mechanisms are largely known and surface molecule expression defining the immunophenotype of each stage are published extensively ^1^.

B-cell development abnormalities or complete blocks are found as a result of monogenic lesions in primary immunodeficiency disorders (PIDD) ^2^. Leukemia and lymphoma of B-cell origin is the most common neoplasia in children, defects in B-cell development and regulation are frequent causes of immune dysregulation diseases in both children and adults, making the B-cell development and function an attractive therapeutic target. As B-cell targeted therapies (e.g. anti-CD20 monoclonal antibodies, anti-CD19 CAR-T) become available and their usage is increasing, iatrogenic B-cell developmental failures are becoming common conditions ^3^.

However, our understanding of the particularities of B-cell developmental abnormalities in those conditions is still incomplete. We currently lack detailed knowledge of the dynamics of additional, non-canonical molecules (new phenotype markers, signaling molecules, therapeutic targets, *in vivo* response to therapy markers). Second, we lack detailed insight into within-a-stage changes, details of transitions or intermediate stages. Third, we lack tools to disclose additional, alternative or non-dominant trajectories potentially present in human patients.

Recent advances in single cell analysis extended the capabilities of clinical flow cytometry beyond 10 parameters, and in another quantum leap forward, spectral ^4^ or mass cytometry ^5^ enabled us to investigate 40 parameters on each cell ^6^. In a proof of principle work of Bendall et al. 2014 ^7^, Wanderlust algorithm was applied to B-cell developmental mass cytometry data, showing assembled progression of markers in a single pathway. We have previously proven that a single 10-color flow cytometry tube is capable of describing the crucial stages of B-cell development and its abnormalities found in PIDD with monogenic lesions in the scope of EuroFlow consortium standardized protocol ^8^. This knowledge is essential, since the inherent assumption of a single-cell trajectory inference is that data contain all markers needed to distinguish all stages and their transition points. Recently, Saelens et al. 2019 ^9^, benchmarked 45 trajectory inference algorithms out of 70 available, concluding that only several would allow for multiple endpoints discovery. Most are built for single-cell RNA data, where the number of cells analyzed is low (10 000) but the number of parameters is high, which contrasts with the mass cytometry dataset, where tens of millions of cells are analyzed with several dozens of parameters. Our objective was to limit the amount of prior information to the starting cell subset, generate all putative random walks and allow for their graphical and user-friendly interrogation and in depth analysis of the selected trajectories.

In the current study, we set out to develop mass cytometry protocol and analytical tools that would allow us to interrogate the B-cell developmental pathways in more detail. We use the 10-color EuroFlow tube as a benchmark. We interrogate the pathways of development leading to terminal cell types expressing either κ light chain or λ light chain across two tissue types.

## Methods

### Sample cohort composition and preparation

Fresh human bone marrow samples (n=3) were obtained from pediatric patients with excluded hematological disease or immunological disorder or (n=1) from fully recovered patient 1 year after successful B-cell precursor leukemia therapy. Only leftover part of the clinical material was used where Informed consent was given. Study was conducted within a project approved by University hospital Motol ethical board. B cells were isolated from the samples using RosetteSep Human B cell Enrichment Cocktail (Stemcell Technologies, Vancouver, BC, Canada) following manufacturer’s instructions. Isolated bone marrow B cells and precursors were cryopreserved in fetal bovine serum containing 10% DMSO in liquid nitrogen. Peripheral B cells were isolated using the same method either from fresh human peripheral blood (n=1) or from buffy coats (n=3), washed with MaxPar Cell Staining Buffer (Standard BioTools, South San Francisco, CA, USA) and used immediately for staining. Bone marrow B cells were thawed for 1 min in 37°C water bath and rested for 30 min in RPMI medium at 37°C in an incubator and washed. Individual samples were barcoded ^10^ with a combination of anti-CD45 and anti-HLA-I metal-tagged antibodies listed in (Supplementary table 1) as described previously ^11^, pooled and further processed in individual tubes.

### Sample staining and acquisition

Metal-tagged antibodies were either purchased (Standard BioTools) or conjugated in-house using Maxpar X8 Antibody Labeling Kit (Standard BioTools) according to manufacturer’s instructions. Antibodies were validated and titrated for the appropriate concentrations and are listed in (Supplementary table 1). The samples were stained as described previously ^12^ and according to the MaxPar Nuclear Antigen Staining with Fresh Fix (Standard BioTools) protocol as described by the manufacturer. Mass cytometry sample acquisition was performed on Helios instrument (Standard BioTools, CyTOF 6.7.1014 software) after preparation according to the manufacturer’s recommendation. Flow cytometry measurement of B-cell precursors was performed exactly as in Wentink et al. 2019 ^8^.

### Data analysis

Acquired samples were exported into FSC format and analyzed manually using sequential bivariate gating in FlowJo (v10.5, FlowJo LLC) software. First, we gated nucleated cells positive for DNA intercalator tagged with 191/193Ir and next the cells positive for particular CD45 and MHC-class I antibody reagent combinations were gated to resolve the barcodes of the individual bone marrow or peripheral blood samples. Next, cell populations for both mass and flow cytometry panels were defined as described previously ^8, 13, 14^, gating strategy shown in Supplementary figure 1 and 2. When markers differently expressed by subsets were sought, we used “population comparison” tool in FlowJo, with probability binning and Cox chi-square statistics.

### Projection with *vaevictis*

For visualization of the mass cytometry data, we used the deep learning-based dimensionality reduction technique using the *vaevictis* model ^15^, one of the autonomous modules integrated in the *tviblindi*. For the projection, the healthy bone marrow (n=4) and healthy peripheral blood (n=4) samples were manually debarcoded and exported as individual FCS files. Next, only cells defined as CD34+ or CD19+ were concatenated into one FCS file and subsequently used for training of the *vaevictis* algorithm, where all panel markers were used for the calculation. Such a trained *vaevictis* model was then applied separately to either the set of bone marrow or the set of peripheral blood cells.

### Trajectory inference in *tviblindi*

For trajectory inference (TI), we used our recent framework called *tviblindi* ^15^, an algorithm integrating several autonomous modules - pseudotime inference, random walk simulations, real-time topological classification using persistence homology, and autoencoder-based 2D visualization using the *vaevictis* model. For the TI, the same concatenated FCS file as for the *vaevictis* projection was used. As a point of origin, stem cells (CD19^-^CD79α^-^TdT^-^CD34^+^) were used. In total, 5000 random walks were probed across the single cell space. Endpoints for further investigation were selected in *tviblindi* graphical user interface (GUI). Topological landmarks were selected in the persistence homology graph in the GUI. Next, walks clustering together were selected on the dendrogram of persistence homology and visually inspected on the *vaevictis* plot. The pseudotime vs. marker line plots were created and exported from the *tviblindi* GUI. For manual analysis of the data in FlowJo, an enhanced FCS file was exported from the *tviblindi* GUI containing all calculated parameters.

## Results

In order to study human B-cell development, we designed a 30-parameter mass cytometry panel allowing for simultaneous measurement of B-cell specific phenotypic surface markers and functional intracellular proteins (Supplementary table 1). We validated the correct assignment of the B-cell precursor subsets by the Euroflow 10-parameter flow cytometry diagnostic panel ^8^. We found a similar distribution of B-cell subsets (gated as in Supplementary Figure 1) measured in four bone marrow samples by mass cytometry and Euroflow flow cytometry, confirming that the mass cytometry panel can describe the basic stages of the B cell development in the bone marrow (Supplementary Figure 3).

Next, we visualized the B-cell precursor subsets using *vaevictis*, a representation learning dimensionality reduction tool built for development visualization. We could observe that the expected main features of the B-cell precursor to mature B-cell development were apparent on four bone marrow and four peripheral blood samples (Figure 1). *Vaevictis* plots of all of the samples individually can be found in Supplementary Figure 4. Using information of all 30 markers, *vaevictis* positioned the mature B-cells adjacent to the B-cell precursors (Figure 1A and 1B), where the aforementioned gated subsets were ordered from the progenitors to mature cell types. Also, the light chain expression highlighted the κ and λ branching (Figure 1C). Progression of the canonical markers (CD34, TdT, CD10, surface IgM (sIgM), κ chain, λ chain, IgD and CD27) in the plot corresponds with the expected course of B cell development (Figure 1C).

**Figure 1.**
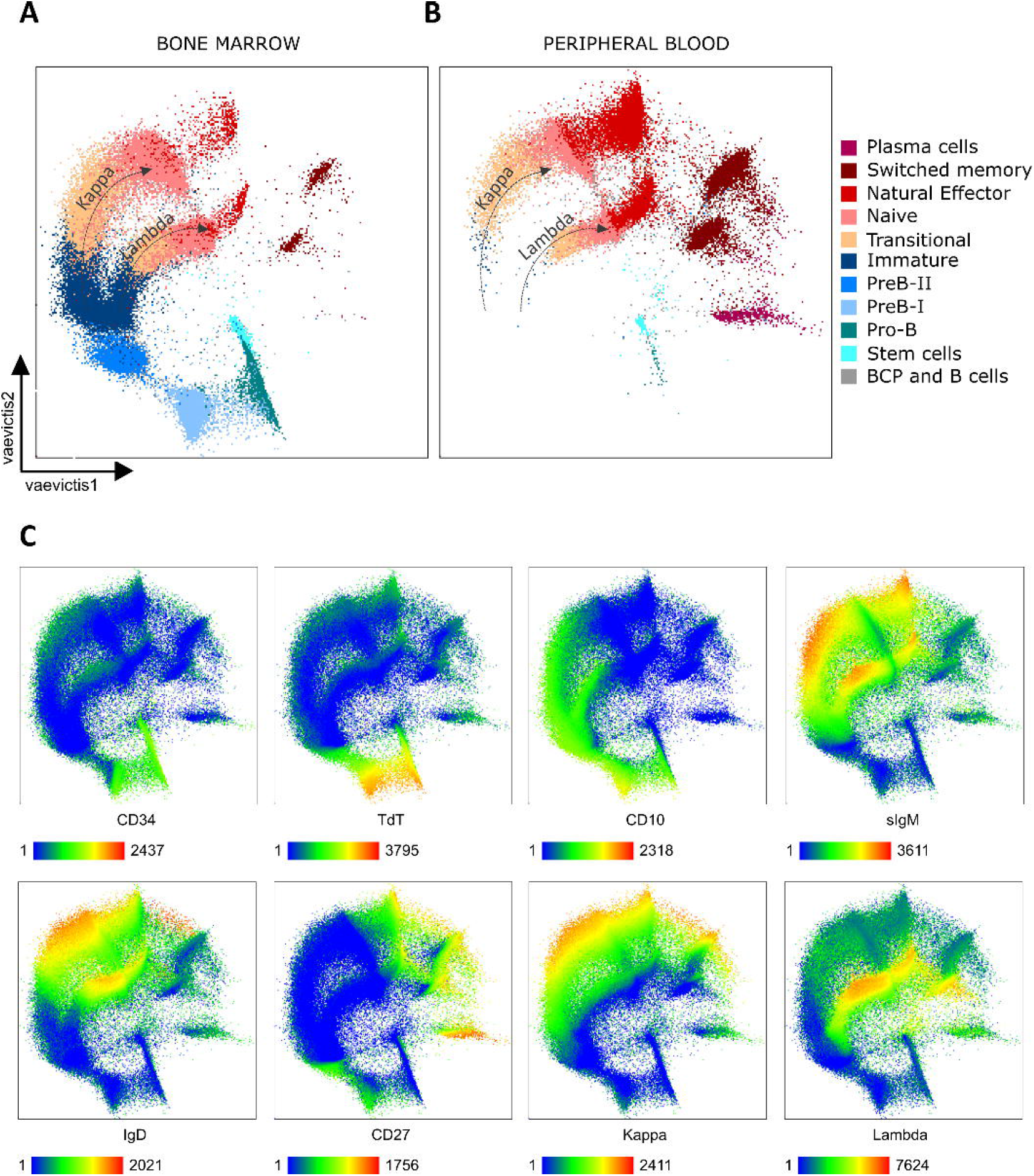
*Vaevictis* dimensionality reduction of BCPs and B cells in bone marrow and peripheral blood. (A) Healthy bone marrow (n=4, concatenated) and (B) peripheral blood (n=4, concatenated) BCPs and B cells with manually gated populations applied to the visualization in color, with annotation and counts of the individual subsets. Dotted arrows highlight kappa and lambda B cells (compare to panel C) (C) Visualization of the entire merged data with the expression of chosen canonical markers using a heatmap color gradient where green represents the lowest expression and red the highest.

Thus, the B-cells and their precursors measured by mass cytometry panel and visualized using *vaevictis* provided bases for interpretation of the putative trajectories of B-cell development.

On the concatenated dataset we selected the developmental point of origin at CD34+ Stem cells. The *tviblindi* algorithm ^15^ was tasked to construct 5000 random walks directed away from the origin (CD34+ Stem cell) with respect to the calculated pseudotime on the nearest neighbor graph (KNNg) of all single cell events. As the KNNg is directed by the pseudotime, endpoints are automatically detected when a random walk reaches a vertex (single-cell event) with no out-going edges. Sixteen endpoints were located in the 6 B-cell subsets corresponding to mature Naive, Natural Effectors and Switched Memory B-cells expressing either κ or λ light chain (Figure 2A). We have selected all endpoints leading to each subset individually. Next, we assembled random walks into different coherent trajectories leading to Natural Effector κ and λ (Figure 2B and C) and to Switched Memory κ and λ (Figure 2D and E) on dendrograms of groups of walks, grouped with respect to the persistent homology classes (Supplementary figure 5 and 6). In parallel we visualized them on the *vaevictis* plot. Since *tviblindi* algorithm and *vaevictis* visualization operate independently in the dataset, we have prioritized abundant walks with particular topology in all dimensions (selected on persistent homology diagram and dendrogram) and those that were transiting through expected cell subsets in a logical sequence (compare Figure 2B, C, D, E to Figure 1A).

**Figure 2.**
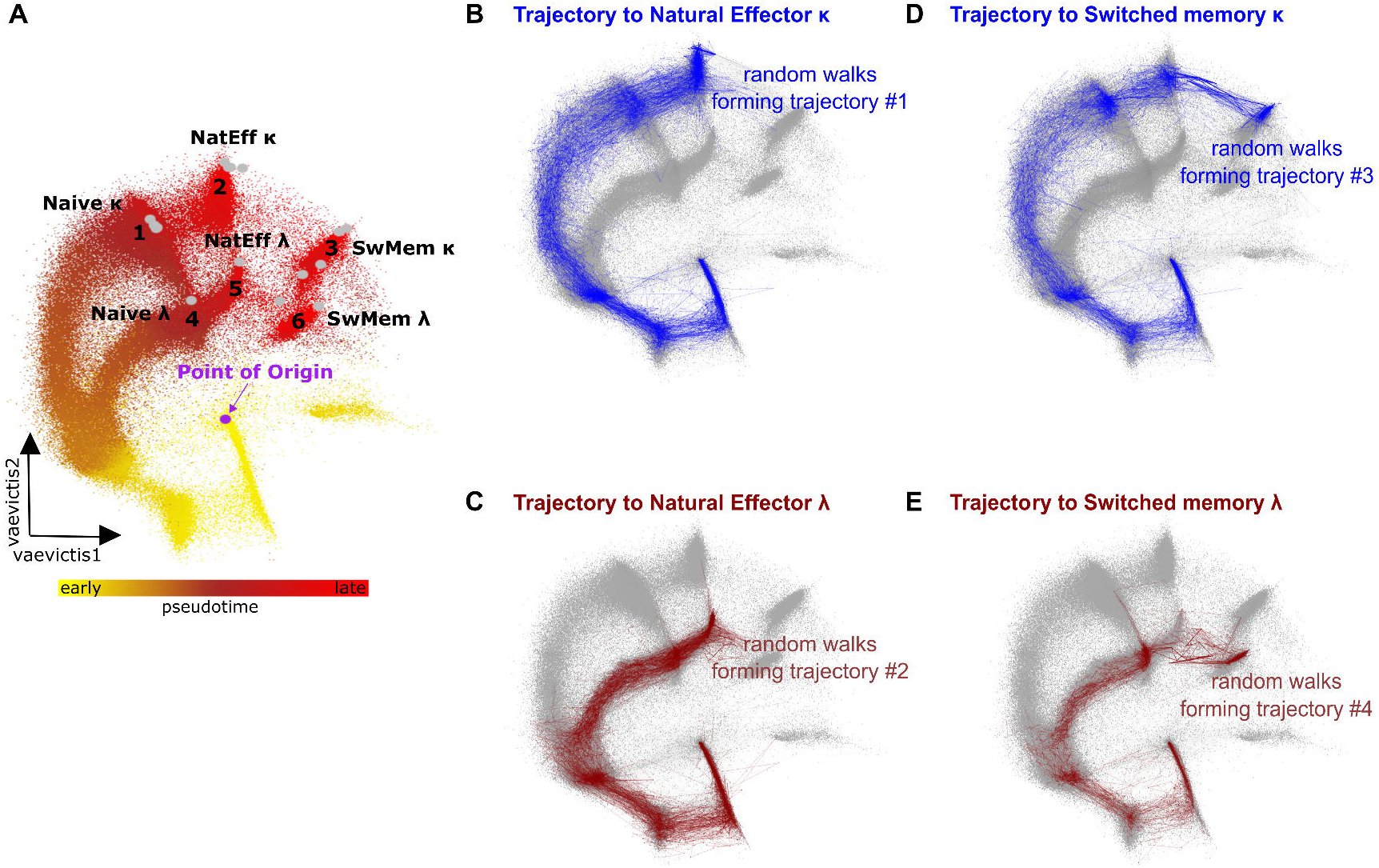
B cell developmental endpoints and trajectories leading to Natural Effector and Switched memory κ and λ B cells constructed by *tviblindi*. (A) Groups of endpoints (1-6) represented as gray dots are located in clusters corresponding to Naïve (1;4), Natural Effector (2;5) and Switched memory (3;6) cells in the vaevictis visualization colored by pseudotime. Yellow color indicates the earliest pseudotime, bright red color indicates the latest pseudotime. The purple dot indicates the Point of origin at gated CD34+ Stem cells. Vaevictis visualization with displayed trajectories to (B) Natural Effector κ and (C) λ B cells and (D) Switched Memory κ And (E) λ B cells constructed by tviblindi (for a selection of trajectory group see (Supplementary figure 5 and 6).

For further analysis, we selected the trajectory leading to Natural Effector κ. We aimed to investigate changes of expression of the markers along the selected developmental trajectory manually. The *tviblindi* interface (GUI, Supplementary Figure 7) allowed us to add all calculated parameters (*vaevictis* 1, *vaevictis* 2, pseudotime, cell assignment to trajectory) and manually investigate the enhanced FCS file for the expression of selected markers along the pseudotime of the selected trajectory (κ, λ and TdT markers shown) (Supplementary Figure 8). Next, we aligned the relative expression values of TdT, CD10 and sIgM in manually gated (as in Supplementary Figure 2) populations of B cell development (Figure 3A) to their expression over the course of pseudotime in the trajectory (Figure 3B). In agreement between manual analysis and pseudotime inference, we found the maximum level of nuclear TdT in ProB/PreB-I stage, CD10 in PreB-I stage and sIgM in Transitional B-cells (Figure 3B), however, the pseudotime plots showed single cell data with all gradual transitions. Thereafter we examined the dynamics of expression of other markers in the early (Stem cells to Pre-BII) (Figure 3C), mid (Pre-BII to Transitionals) (Figure 3D), and late (Transitionals to Natural Effectors) (Figure 3E) phases of B-cell development. See Supplementary Figure 9 for a continuation of the expression to the Switched Memory subset.

**Figure 3.**
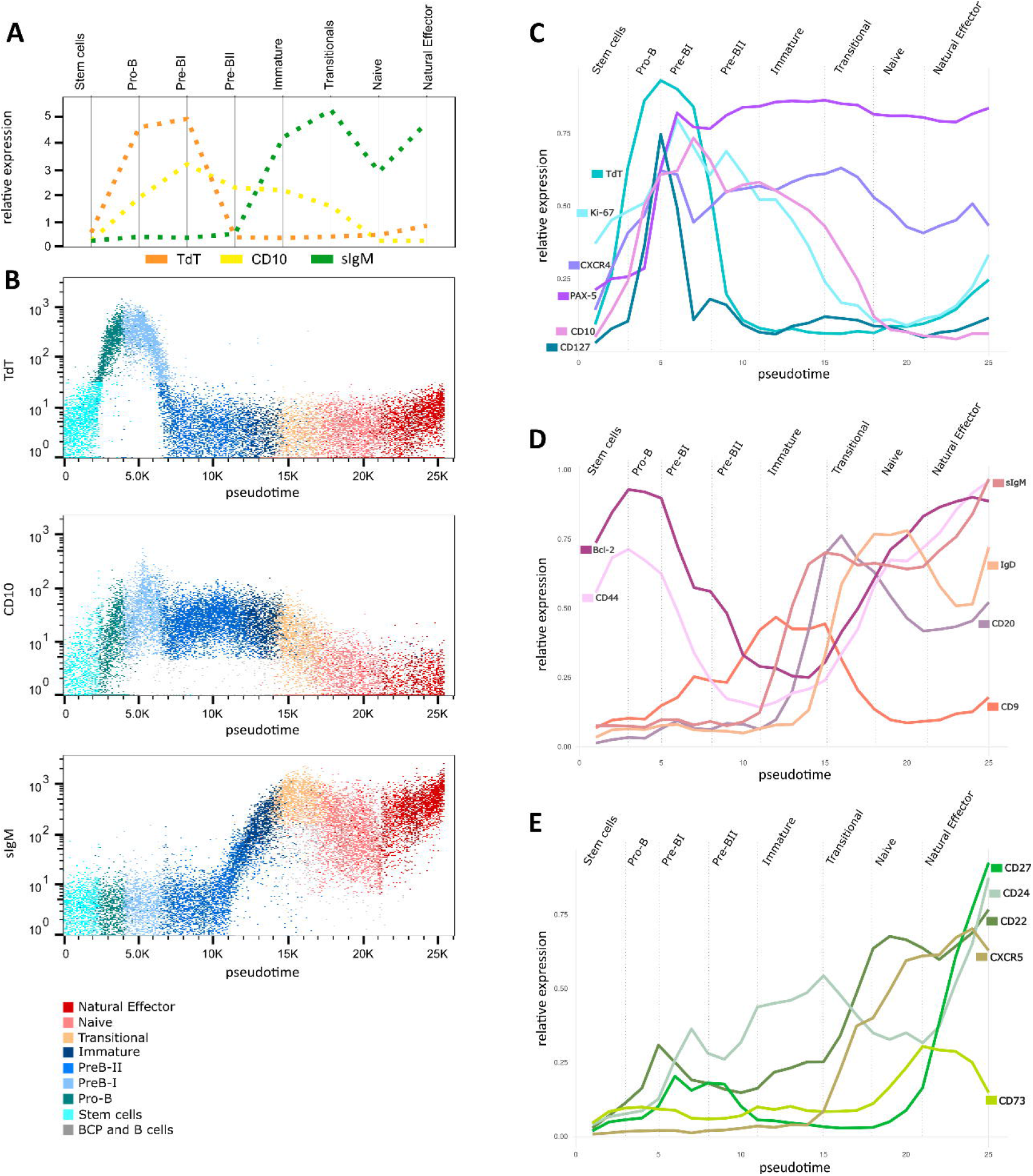
Detailed analysis of the trajectory leading to Natural Effector κ cells. (A) Median expression of TdT (orange), CD10 (yellow) and sIgM (green) from manually gated populations correlate with (B) the expression of TdT, CD10 and sIgM on the pseudotime vs. marker dot plots with manually gated populations overlaid in color. Pseudotime line plots showing the average expression of markers upregulated in the early (C), mid (D) and late (E) phase of the development, manually annotated by the gated stages.

TdT expression in the Pro-B cells was followed by the CD127 (IL-7R), CXCR4, PAX-5, Ki-67 and CD10 expression at Pro-B to Pre-BI transition. Notably, CD127 raised and declined before Ki-67 peaked in Pre-BI while a second (smaller) CD127 peak followed by Ki-67 was seen in Pre-BII, in line with the reported role of IL-7 signaling inducing proliferation in humans ^16^. Similarly, PAX-5 peak follows the CD127 peak (Figure 3C). The Bcl-2 and CD44 present bimodal expression, peaking at Pro-B stage first and again in mature stages in the peripheral blood (Figure 3D). CD9 peaks within the Immature stage, followed by sIgM, while CD20 and IgD peak at Transitional B-cell stage (Figure 3D). Finally, the CD22 and CXCR5 increase to their peaks at Naive and Effector stage (Figure 3E), respectively, followed by CD73, which is down modulated in the Natural Effector cells and upregulated again in the Switched memory B cells (Supplementary figure 9C). CD27 is known as a B-cell memory marker, but in fact has also bi-modal expression with a first peak at Pre-BI and Pre-BII stages and second peak at memory stages. The expression of CD24 is first elevated in the Pro-B to the Transitional stage only to reach its highest level in Natural Effectors (Figure 3E).

Analyzing the two compartments separately (Figure 4A, 4B), we could see that Transitional, Naive and Natural Effector cells were present in both the bone marrow and the peripheral blood. Their pseudotime position was slightly different suggesting there is a phenotypic difference. Indeed, we found higher expression of CXCR4 and lower expression of CXCR5 in the subsets in the bone marrow compartment (Figure 4C). The comparison of the Switched Memory subsets is shown in Supplementary Figure 10.

**Figure 4.**
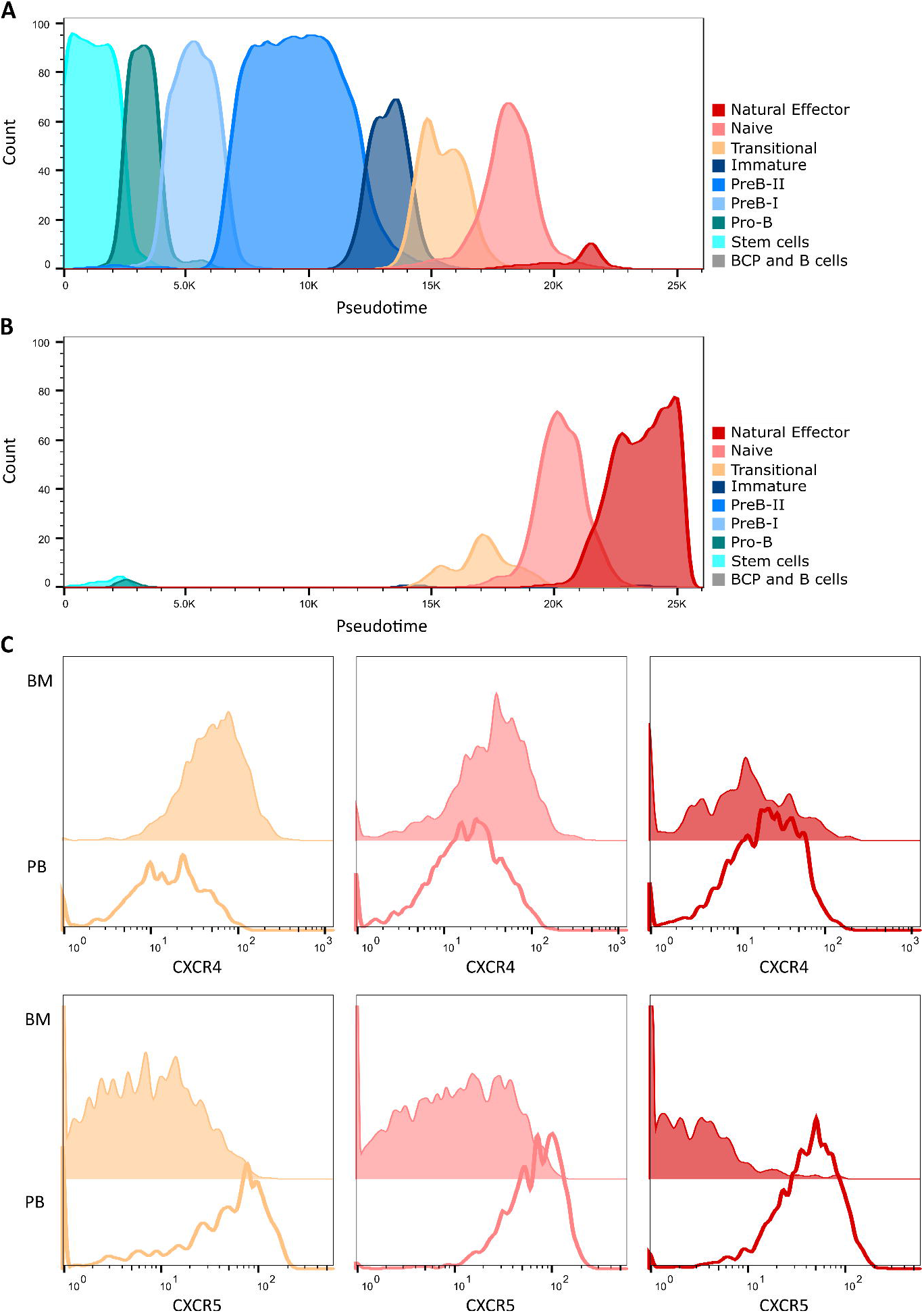
Contribution of the BCP and B-cell stages to the bone marrow and peripheral blood compartments. Individual populations are shown as histograms in the data set divided into (A) bone marrow (n=4, concatenated) and (B) peripheral blood (n=4, concatenated). (C) Differential expression of the markers CXCR4 (top row) and CXCR5 (bottom row) in the populations which are present in both of the compartments. Solid line histograms indicate subsets present in the peripheral blood (PB). Filled histograms indicate subsets present in the bone marrow (BM).

Finally, we set out to find and describe the branching points of discovered trajectories. We easily identified the position of the branching of trajectories to Naive κ and λ endpoints when plotting the κ light chain versus pseudotime (Figure 5A). As expected, the branching point was located at the Pre-BII to Immature transition on a *vaevictis* projection (Figure 5B). Following analogous principle, we investigated the branching point in the development of cells into natural effector and Switched Memory cells (the selection of random walks on dendrograms can be seen in Supplementary figure 11). After selecting the two trajectories based on the respective endpoints, we sought differentially expressed markers by the endpoint subsets. CD73 was the most different of all markers measured. Plotting the two trajectories versus CD73, we found a branching point at a trajectory segment were CD73 started to increase on the way to the switched memory B-cells (Figure 5C). The branching point was topologically located at the Naïve B-cells (Figure 5D). While examining the trajectory to natural effector B-cell endpoint, we noticed that CD73 was heterogenous and investigating of the dendrogram of walks we found two similar trajectories, one devoid of CD73 expression and a second with only transient increase of CD73 expression (Supplementary figure 12). The expression of CD73 was preceded by CXCR5, a germinal center homing marker.

**Figure 5.**
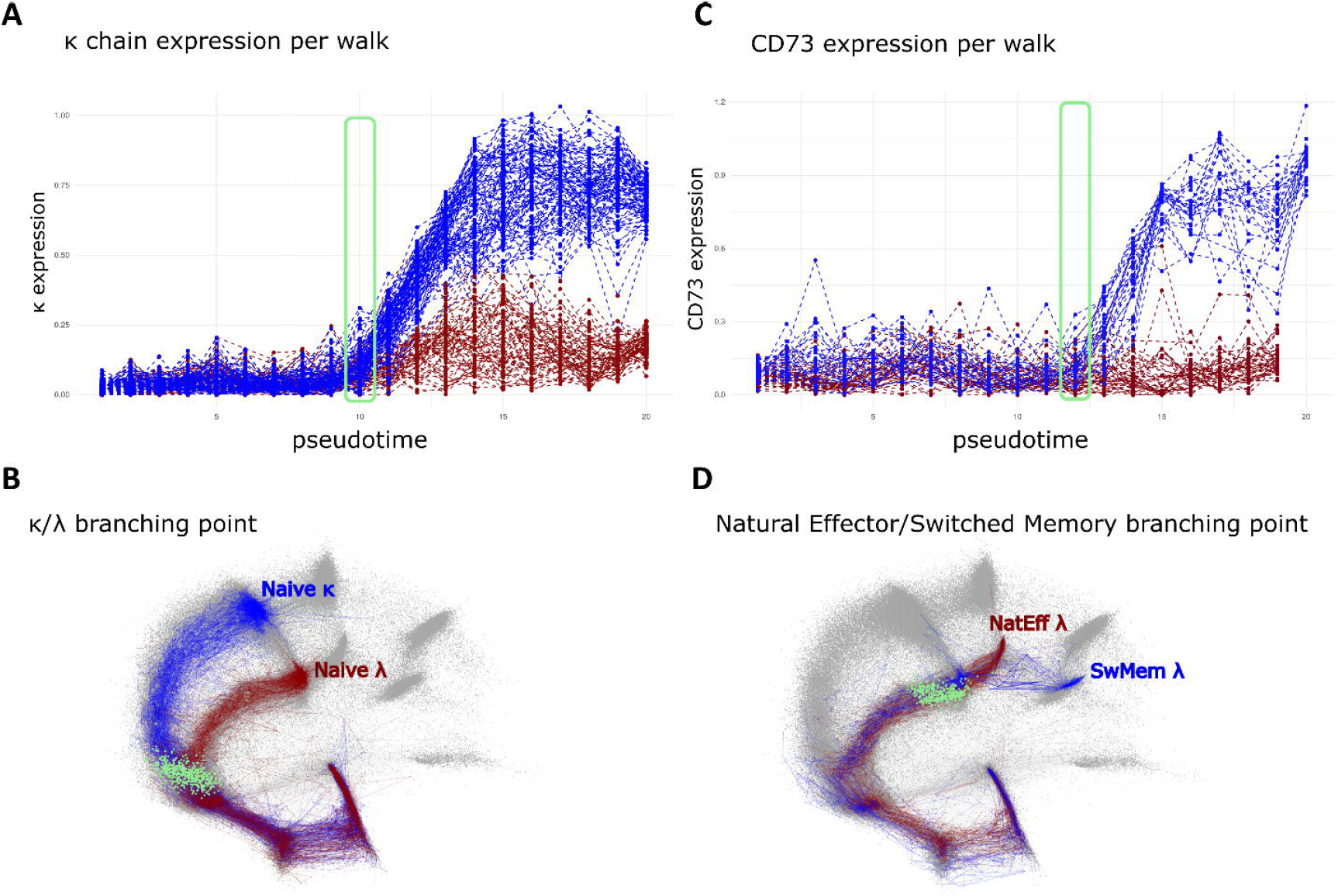
Identification of branching points in trajectories leading to the selected endpoints. (A) Pseudotime line plot showing expression of the κ chain for the selected trajectories leading to Naive κ (blue) and Naive λ (red) ends. The green rectangle indicates the branching point of trajectories and is projected as green dots in the *vaevictis* plot. (B) *Vaevictis* plot showing trajectories to Naive κ (blue) and Naïve λ (red) subsets with projection of cells (green) located in the branching point. (C) Pseudotime line plot showing the expression of the CD73 for the selected trajectories leading to Switched memory λ (blue) and Natural Effector λ (red) ends. The green rectangle indicates the branching point of trajectories and is projected as green dots in the *vaevictis* plot. (D) *Vaevictis* plot showing trajectories to Natural Effector λ (red) and Switched memory λ (blue) endpoints with projection of cells (green) located in the branching point.

## Conclusions and discussion

We presented a single cell analysis solution for interrogation of B-cell developmental trajectories on multiple samples of relevant tissues (bone marrow and peripheral blood). We designed and validated a mass cytometry panel capable of evaluating 30-markers plus 5 sample barcodes. We compared its performance to a benchmark of EuroFlow 10-color cytometry assay. We showed a practical, feasible and scalable *vaevictis* projection calculation based on deep learning.

This tool is designed to create a continuous representation of the data rather than isolated clusters allowing a clear interpretation of the dynamics in the data (as compared of other currently used methods t-SNE ^17^ or UMAP ^18^. Due to the naïve importance sampling, numerically dominant populations are not overrepresented in the 2D plot and the running time is basically insensitive to the size of the original dataset. The deep learning architecture allows for direct reuses of the trained representation on a newly acquired sample (if performed in a standardized manner).

Thus, uniquely, samples of different donors (affected and unaffected) and of multiple tissues (central and peripheral) can be probed with thousands of putative pathways, that are defined only by a starting point, markers used and the overall definition of cells belonging to the pool of relevant cell type (here stem cells and B-cell lineage).

All trajectories found are visually presented for interrogation, diverse terminal ends can be selected and individual trajectories are assembled into relevant pathways for further exploration. Recent mathematical apparatus based on persistence homology calculations is used to quantitatively describe similarities of trajectories that can be assembled together.

Notably, trajectories found in our dataset correspond to the known theory of B-cell development, they logically transit from the central organ of hematopoiesis (bone marrow) to the periphery (blood). When dissected in detail, they show expected sequences of canonical markers, but add detail to the transition points and provide dynamic information about the expression of markers within known stages. For example, CD127 peaking before PAX-5 is in line with a recent study showing an important role of CD127 (IL-7RA) signaling in promoting PAX-5 expression ^16^. While CD27 is conventionally used for phenotypic description of memory B cell subsets in the periphery, we show its upregulation also in the Pre-B stages, as shown earlier by Vaskova et al. 2008 ^19^. The transient downregulation of CXCR4 and simultaneous CD9 upregulation which we see within the Pre-BI stage is in line with Leung et al. 2011 ^20^, who observed that CD9 levels are enhanced after SDF-1 stimulation suggesting that CD9 plays a role in the SDF-1/CXCR4 axis known to be essential in HSC/progenitor homing. The expression profile of CD24 along the calculated pseudotime follows the experimental findings of studies ^21, 22^ showing the highest peak of expression in Transitional B cells (followed by a decrease in Naive cells) and the second in memory B cells. While the mature B-cell stages were immunophenotypically similar in the bone marrow and peripheral blood (found in the same regions on *vaevictis* plot), we could find quantitative difference in the CXCR4 and CXCR5 expression, known homing receptors ^23, 24^. Our approach allowed us to investigate multiple developmental endpoints resulting from trajectories’ branching. The CD73, a known marker of switched memory B-cells ^25^ gradually increased until the switched memory B-cell stage, while it remained negative or only transiently increased towards the natural effector B-cell stage. While the branching point was found at naïve B-cell stage, the heterogenous expression of CD73 together with CXCR5 expression suggested that there are still alternative trajectories among the natural effector B-cells. One extrafollicular trajectory seems defined by the absence of CXCR5 and CD73 (CXCR5 and CD73 negative), while a second trajectory, defined by a transient expression of CXCR5, may describe cells that passage transiently in the germinal center. Indeed, the distinction between extrafollicular B cells and natural effector-B cells is still unclear ^26^, and our analysis can provide insights on marker definition to dissect distinct cell fates.

Unlike the so far published algorithms that oversimplify the trajectory inference showing single dominant trajectory or two trajectories with a single branching point (e.g. Wishbone ^27^), we could analyze multiple branching points and bring quantitative expression information as well as topological information about the branching point. The theoretical limitation is the number of investigated markers and choice of tissues and samples. This can be overcome by using *tviblindi* on a single-cell RNASeq or better yet CITE-Seq dataset combining the protein markers with gene expression and enriching the mass cytometry panel in the next iteration. We can generalize, that tviblindi algorithm can reliably show the sequence of expression of surface markers as well as nuclear transcription factors. We could anticipate that our mass cytometry panel and *tviblindi* could be used to compare healthy reference with affected (intrinsic or extrinsic factors) development to discover alternative pathways, new branching points or alternative endpoints. These deviations could potentially disclose targetable processes for therapy of PIDD and/or B-cell neoplasia.

## Acknowledgements

We wish to thank to Daniel Thurner, Pavla Luknarova and Katerina Rejlova for their superb technical assistance. TK and JS were supported by grant 23-05561S from The Czech Science Foundation. TK, DK and JS were funded by the European Union - Next Generation EU (Czech Recovery Plan) - Project National Cancer Research Institute LX22NPO5102. DN was funded by the FWO (Fonds Wetenschappelijk Onderzoek) and conducted as part of the FWO Strategic Basic research project 1S40421N.

