## Supplementary data for "Tviblindi algorithm identifies branching developmental trajectories of human B cell development"

### Supplementary figures

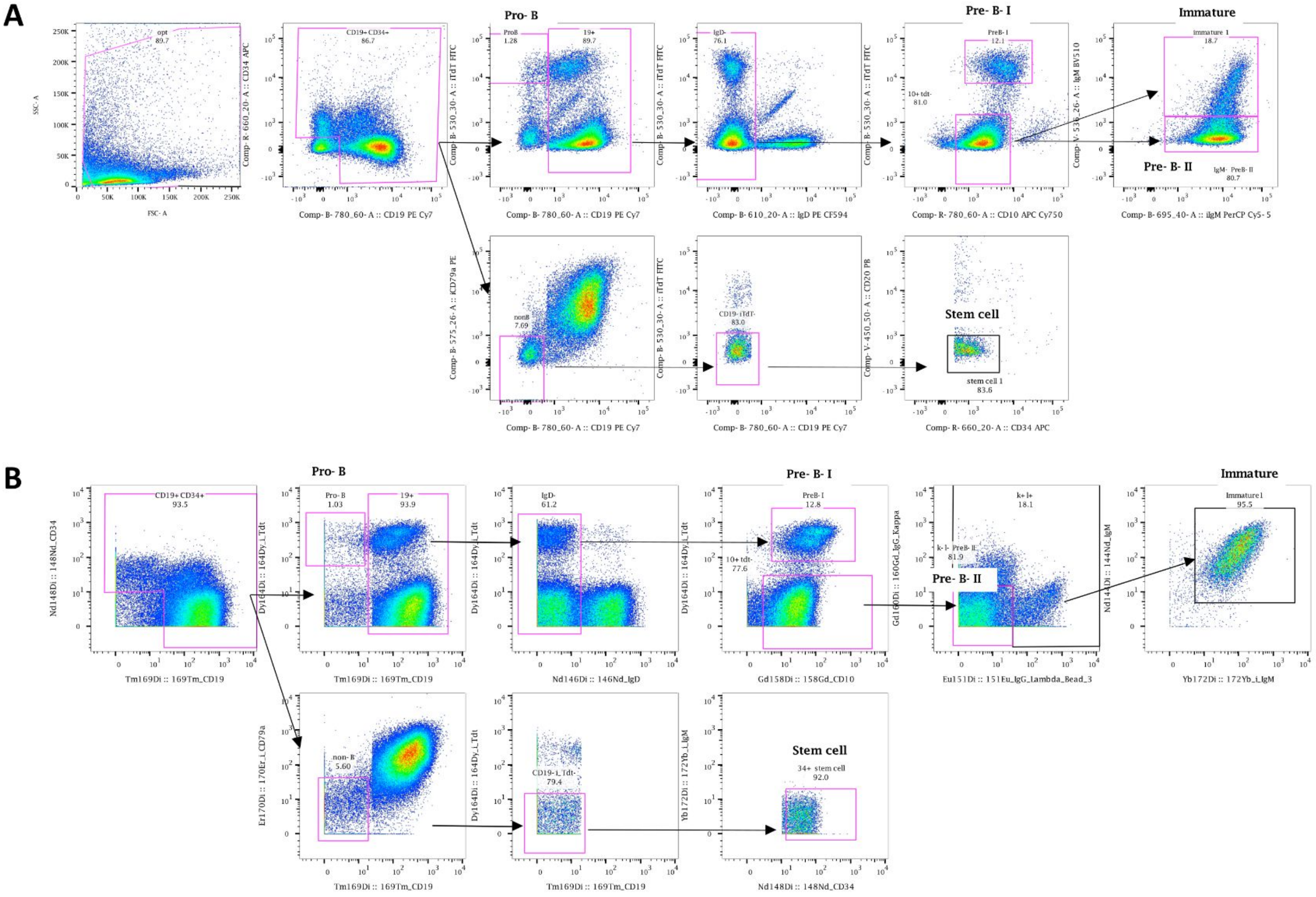


**Supplementary Figure 1. Gating strategy for the flow cytometry (A) and CyTOF (B) data**. Representative results from the same healthy bone marrow sample measured in parallel.

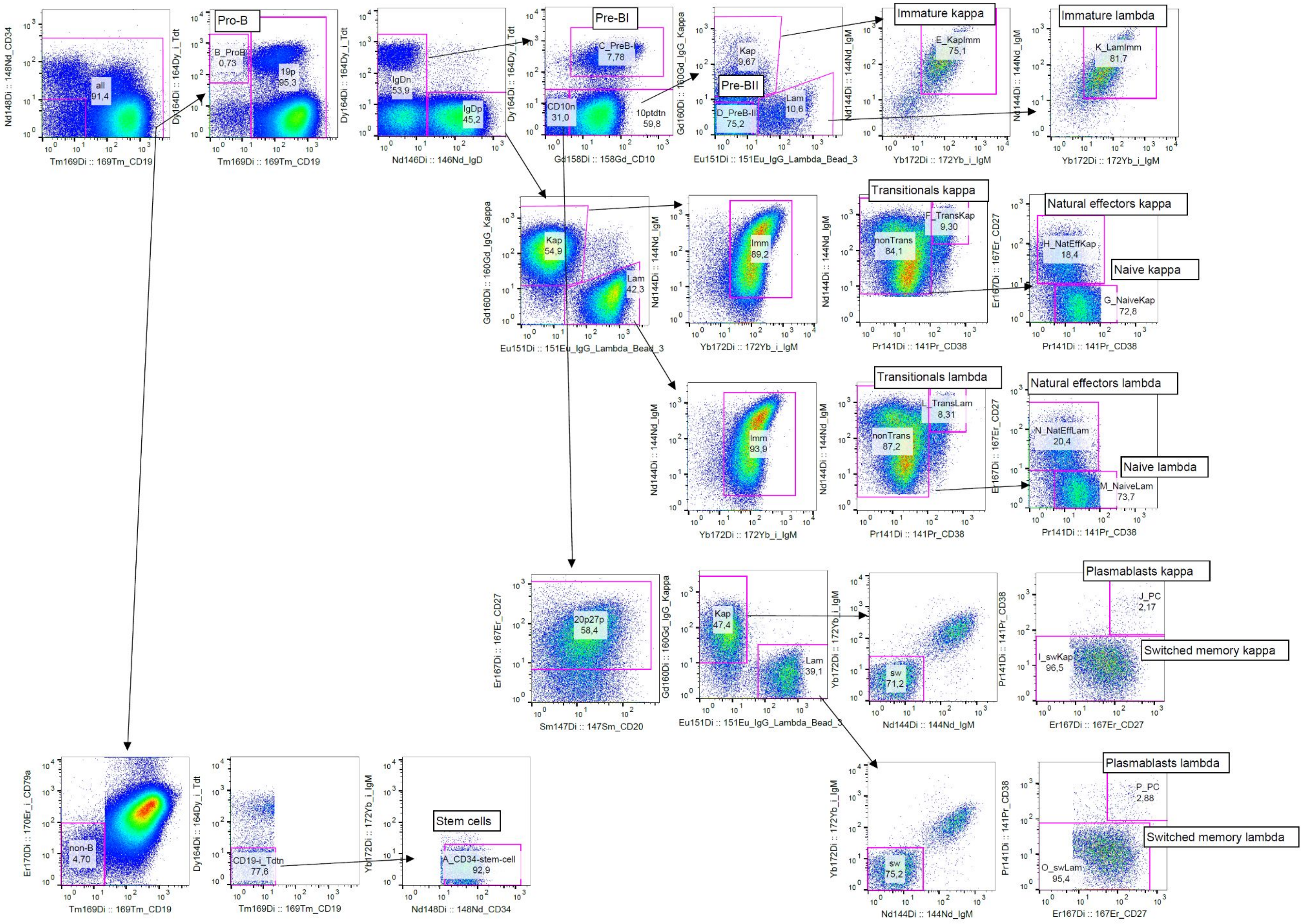


**Supplementary figure 2**. Gating strategy for the CyTOF data from healthy donor bone marrow and peripheral blood samples.

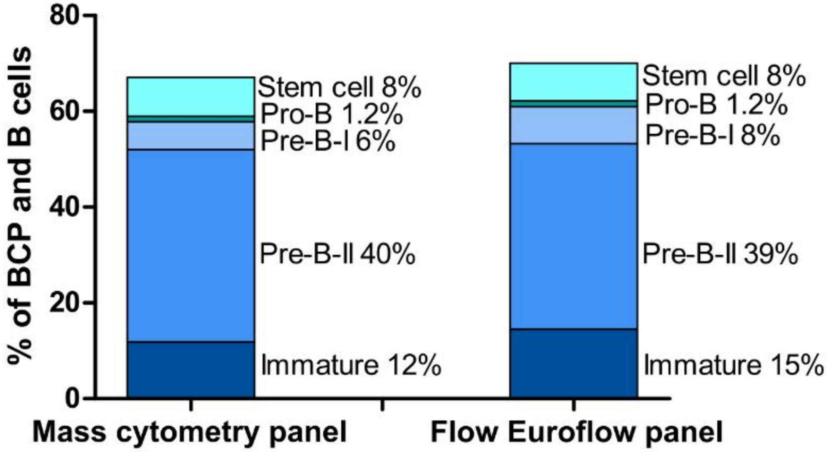


**Supplementary Figure 3. Distribution of B-cell precursor populations correlates between the novel mass cytometry panel (left) and previously validated flow Euroflow panel (right)**. Colors indicate individual populations with percentage representation within both of the panels. Stem cells were defined as CD19-CD79α-TdT-CD34+, Pro-B cells as CD19-TdT+CD34+, Pre-BI cells as CD19+TdT+CD34+, Pre-BII cells as CD19+CD10+iIgM+IgM- and Immature cells as CD19+CD10+iIgM+IgM+. Results from 4 different healthy bone marrow samples measured in parallel.

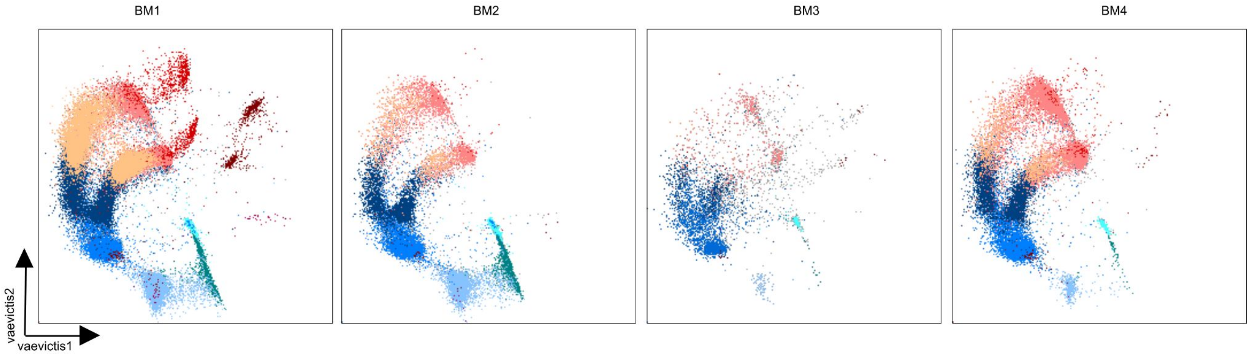


| | Subset Name | Count |
|---|---|---|
| | Plasmablasts | 55,0 |
| | Switched memory | 713 |
| | Natural Effector | 1764 |
| | Naive | 41570 |
| | Transitionals | 9364 |
| | Immature | 9817 |
| | PreB-II | 46390 |
| | PreB-I | 9345 |
| | Pro-B | 1327 |
| | Stem cells | 4825 |
| | BM1 | 135926 |

| | Subset Name | Count |
|---|---|---|
| | Plasmablasts | 8,00 |
| | Switched memory | 24,0 |
| | Natural Effector | 85,0 |
| | Naive | 3328 |
| | Transitionals | 762 |
| | Immature | 3520 |
| | PreB-II | 24082 |
| | PreB-I | 6513 |
| | Pro-B | 1610 |
| | Stem cells | 6218 |
| | BM2 | 59672 |

| | Subset Name | Count |
|---|---|---|
| | Plasmablasts | 1,00 |
| | Switched memory | 19,0 |
| | Natural Effector | 24,0 |
| | Naive | 558 |
| | Transitionals | 30,0 |
| | Immature | 662 |
| | PreB-II | 2132 |
| | PreB-I | 83,0 |
| | Pro-B | 16,0 |
| | Stem cells | 839 |
| | BM3 | 6803 |

| | Subset Name | Count |
|---|---|---|
| | Plasmablasts | 5,00 |
| | Switched memory | 71,0 |
| | Natural Effector | 305 |
| | Naive | 14231 |
| | Transitionals | 1001 |
| | Immature | 5502 |
| | PreB-II | 37954 |
| | PreB-I | 536 |
| | Pro-B | 81,0 |
| | Stem cells | 672 |
| | BM4 | 69935 |

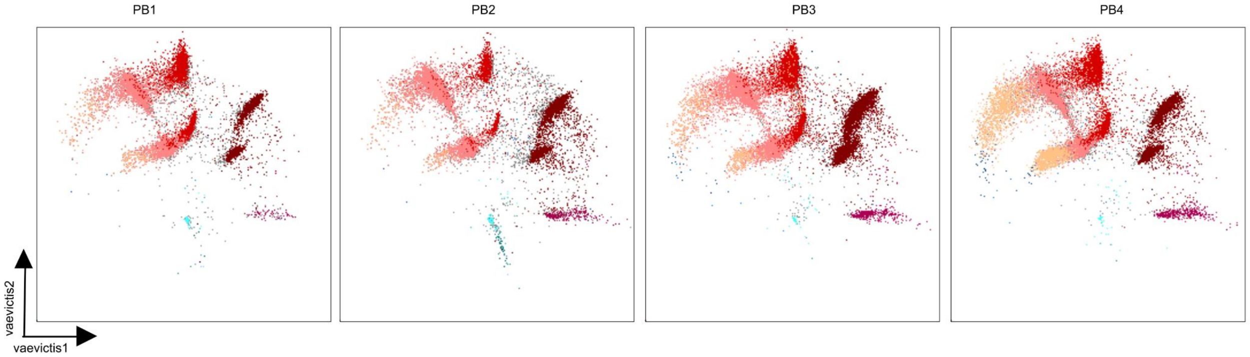


| | Subset Name | Count |
|---|---|---|
| | Plasmablasts | 69,0 |
| | Switched memory | 2290 |
| | Natural Effector | 3445 |
| | Naive | 7955 |
| | Transitionals | 327 |
| | Immature | 6,00 |
| | PreB-II | 7,00 |
| | PreB-I | 5,00 |
| | Pro-B | 6,00 |
| | Stem cells | 46,0 |
| | PB1 | 19000 |

| | Subset Name | Count |
|---|---|---|
| | Plasmablasts | 231 |
| | Switched memory | 4518 |
| | Natural Effector | 1368 |
| | Naive | 9624 |
| | Transitionals | 234 |
| | Immature | 11,0 |
| | PreB-II | 31,0 |
| | PreB-I | 4,00 |
| | Pro-B | 42,0 |
| | Stem cell | 110 |
| | PB2 | 23429 |

| | Subset Name | Count |
|---|---|---|
| | Plasmablasts | 448 |
| | Switched memory | 14881 |
| | Natural Effector | 18903 |
| | Naive | 10895 |
| | Transitionals | 734 |
| | Immature | 81,0 |
| | PreB-II | 67,0 |
| | PreB-I | 2,00 |
| | Pro-B | 1,00 |
| | Stem cells | 53,0 |
| | PB3 | 59372 |

| | Subset Name | Count |
|---|---|---|
| | Plasmablasts | 422 |
| | Switched memory | 5823 |
| | Natural Effector | 5317 |
| | Naive | 48406 |
| | Transitionals | 2556 |
| | Immature1 | 75,0 |
| | PreB-II | 32,0 |
| | PreB-I | 3,00 |
| | Pro-B | 3,00 |
| | Stem cells | 47,0 |
| | PB4 | 81686 |

**Supplementary Figure 4. Vaevictis dimensionality reduction of BCPs and B cells in bone marrow and peripheral blood for the individual samples**. Visualization of the four healthy donor bone marrow samples (top row, BM1-BM4) and the four healthy donor peripheral blood samples (bottom row, PB1-PB4) with manually gated populations applied to the graph in color, with annotation and counts of the individual subsets.

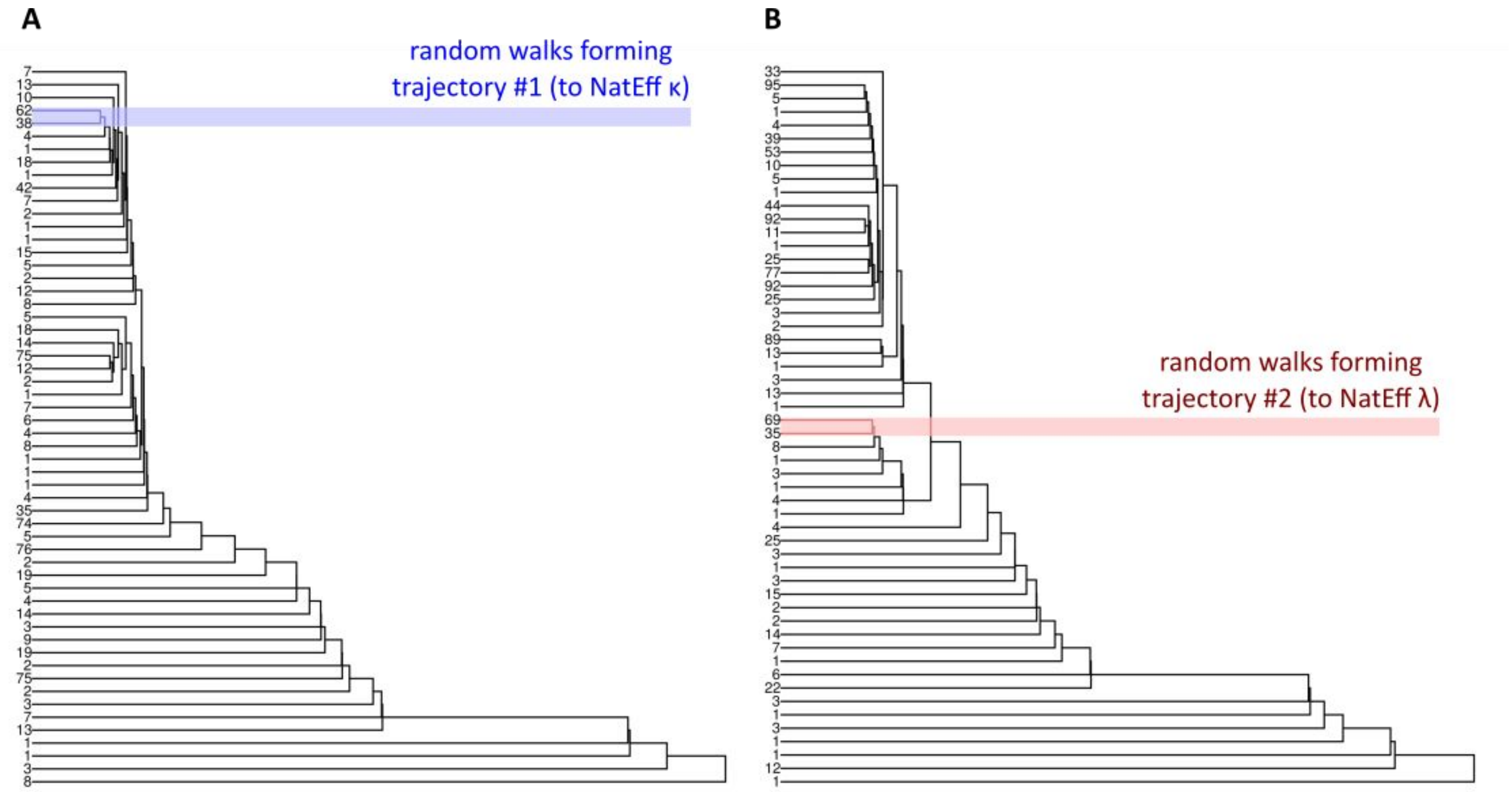


**Supplementary Figure 5. Hierarchical clustering dendrogram for Natural Effector κ and λ developmental trajectories**. Dendrograms show clustering of all random walks leading to developmental endpoints located in the (A) Natural Effector κ and (B) Natural Effector λ clusters. Leaves represent the groups of random walks with similar topology. In blue is highlighted the selected group of walks representing developmental trajectory to Natural Effector κ cells. In red is highlighted the group of walks representing developmental trajectory to Natural Effector λ cells.

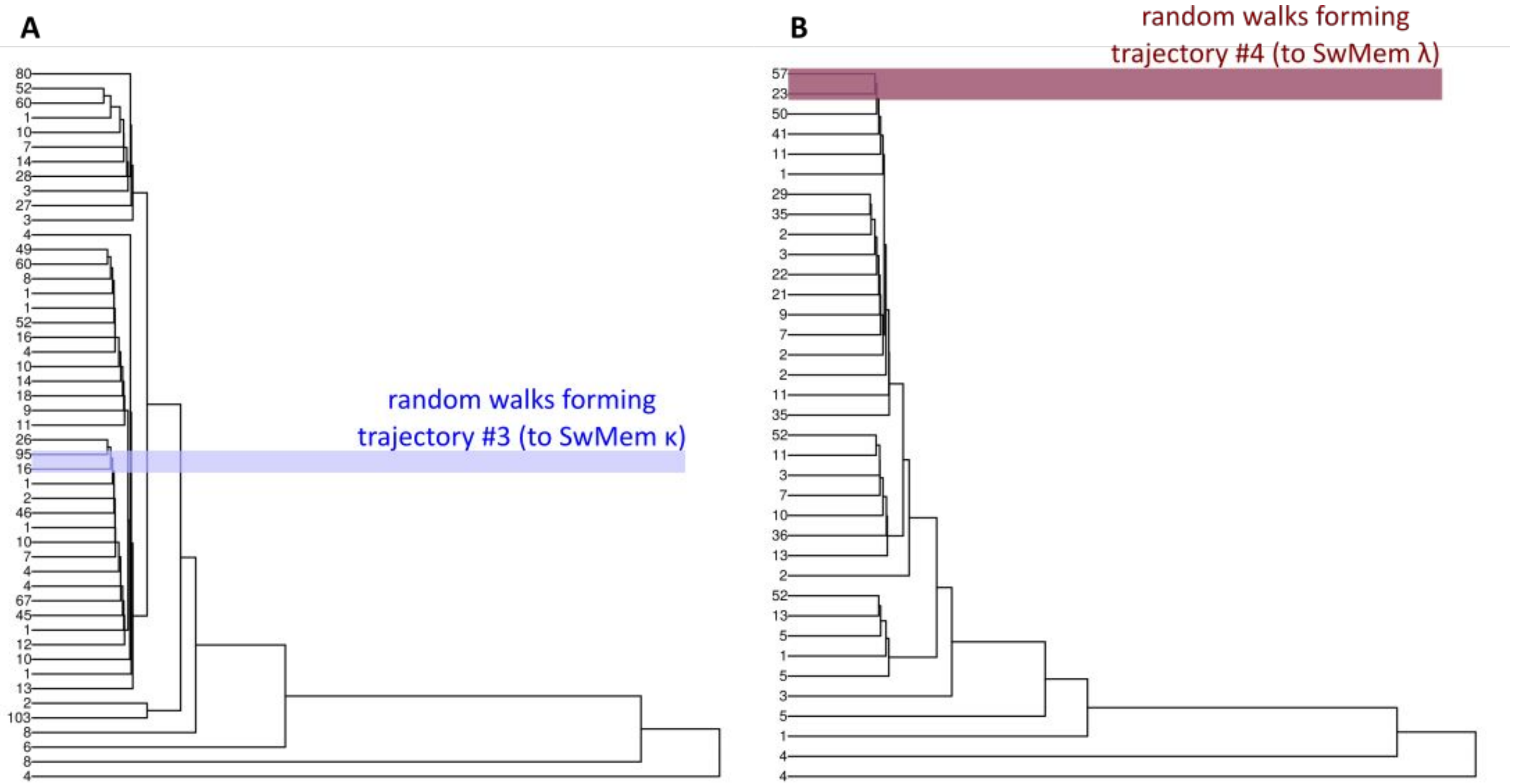


**Supplementary Figure 6. Hierarchical clustering dendrogram for Switched Memory κ and λ developmental trajectories**. Dendrograms show clustering of all random walks leading to developmental endpoints located in the (A) Switched Memory κ and (B) Switched Memory λ clusters. Leaves represent the random walks and their abundance. In blue is highlighted the selected group of walks representing developmental trajectory to Switched Memory κ cells (A) and Switched Memory λ cells (B).

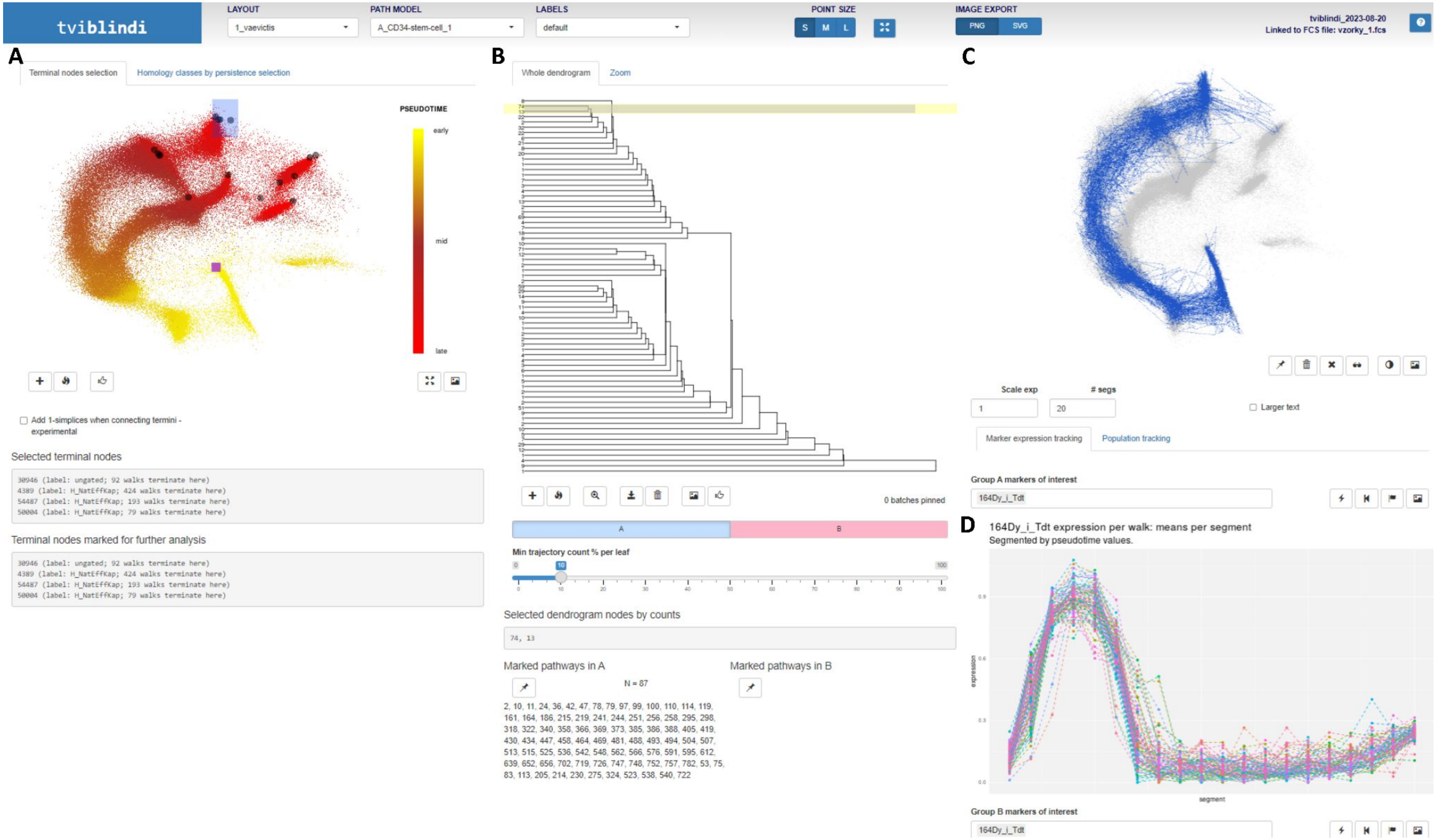


**Supplementary Figure 7. tviblindi graphical user interface (GUI).** (A) vaevictis plot with selected terminal endpoints located in the Natural Effector κ cluster, (B) hierarchical clustering dendrogram with selection of random walks forming the trajectory leading to Natural Effector κcluster, (C) vaevictis plot with random walks selected in (B), (D) lineplot with the expression of TdT along pseudotime of the selected trajectory.

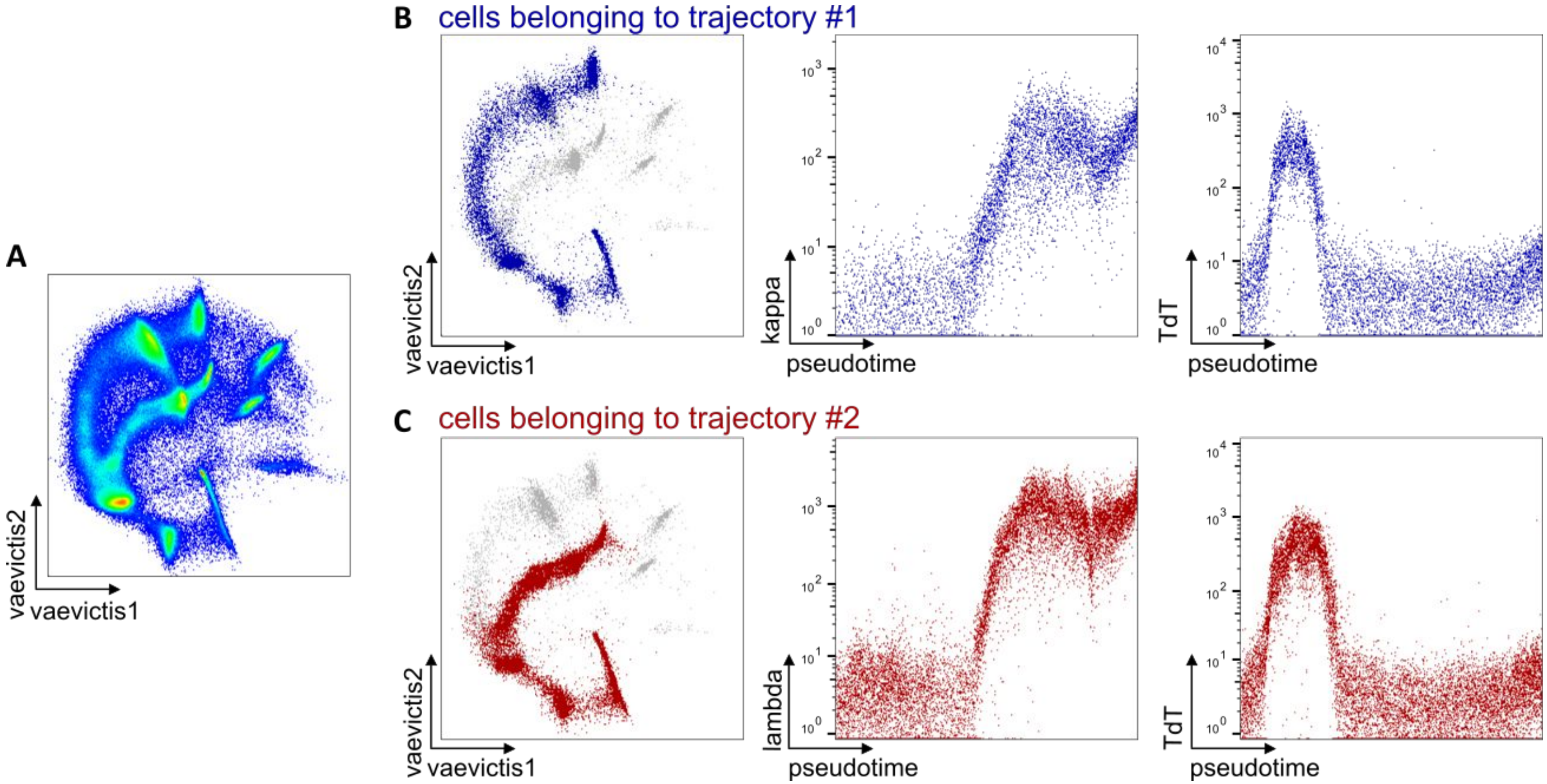


**Supplementary Figure 8. Manual analysis of trajectories constructed by tviblindi (at the cellular level?) using enhanced FCS file**. Vaevictis visualization of (A) the entire data set and (B, C) with colored cells belonging to the selected trajectory ending at the Natural Effector κ (blue) and λ (red) subsets in FlowJo. Expression of κ, λ and TdT markers along pseudotime.

**Supplementary Figure 9. Detailed analysis of the trajectory leading to Switched memory κ cells**. Pseudotime line plots showing the average expression of markers upregulated in the early (A), mid (B) and late (C) phase of the development.

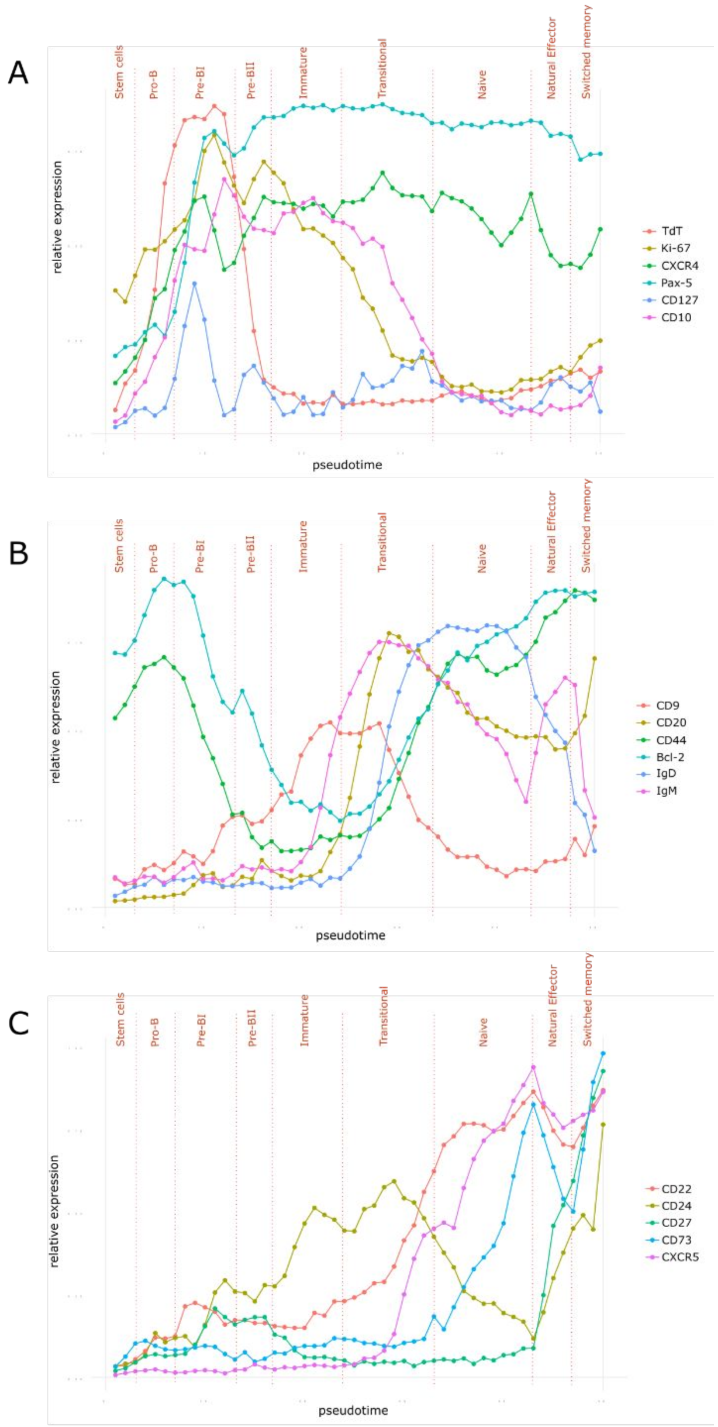

**Supplementary Figure 10.** Differential expression of the markers CXCR4 (A) and CXCR5 (B) in Switched memory cells present in both bone marrow (BM) and peripheral blood (PB) compartments. Solid line histograms indicate subsets present in the PB. Filled histograms indicate subsets present in the BM.

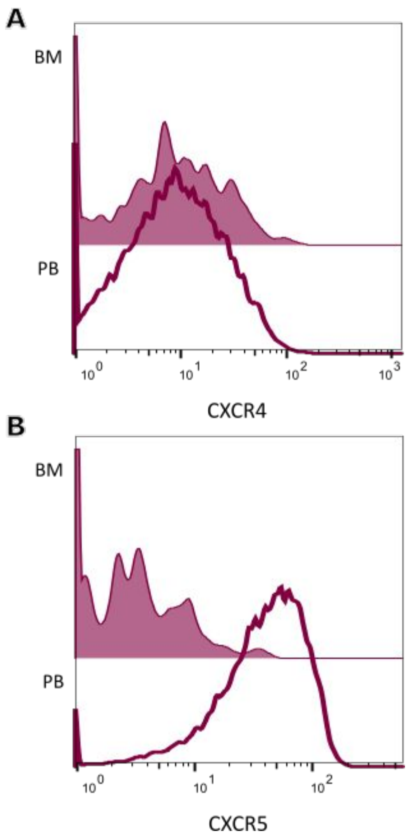

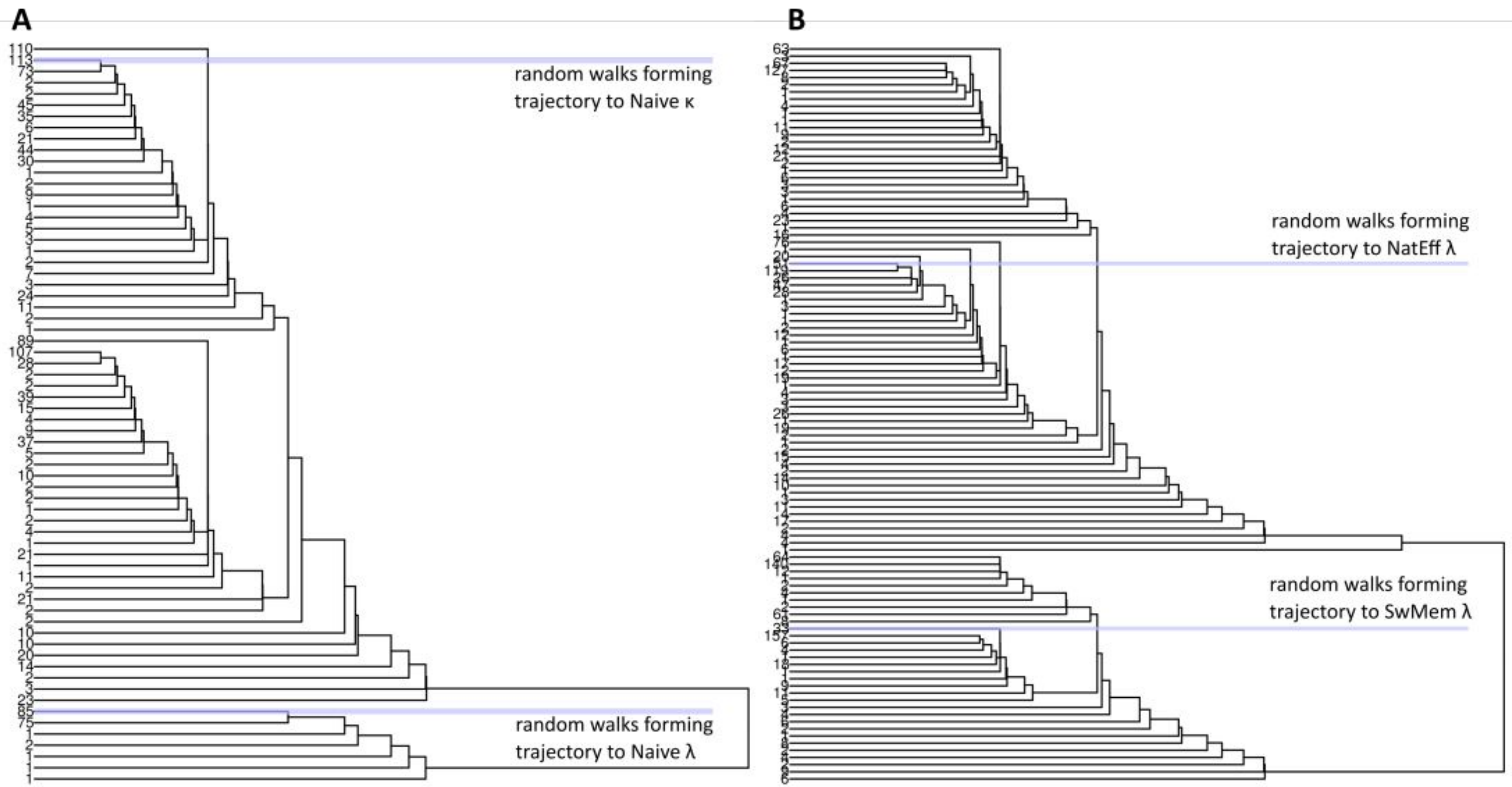


**Supplementary Figure 11. Hierarchical clustering dendrograms.** For the developmental branching of (A) Naive κ and λ cells and (B) Natural Effector λ and Switched memory λ cells. In blue are highlighted the groups of random walks representing the developmental trajectories to (A) Naive κ and λ cells and (B) Natural Effector λ and Switched memory λ cells.

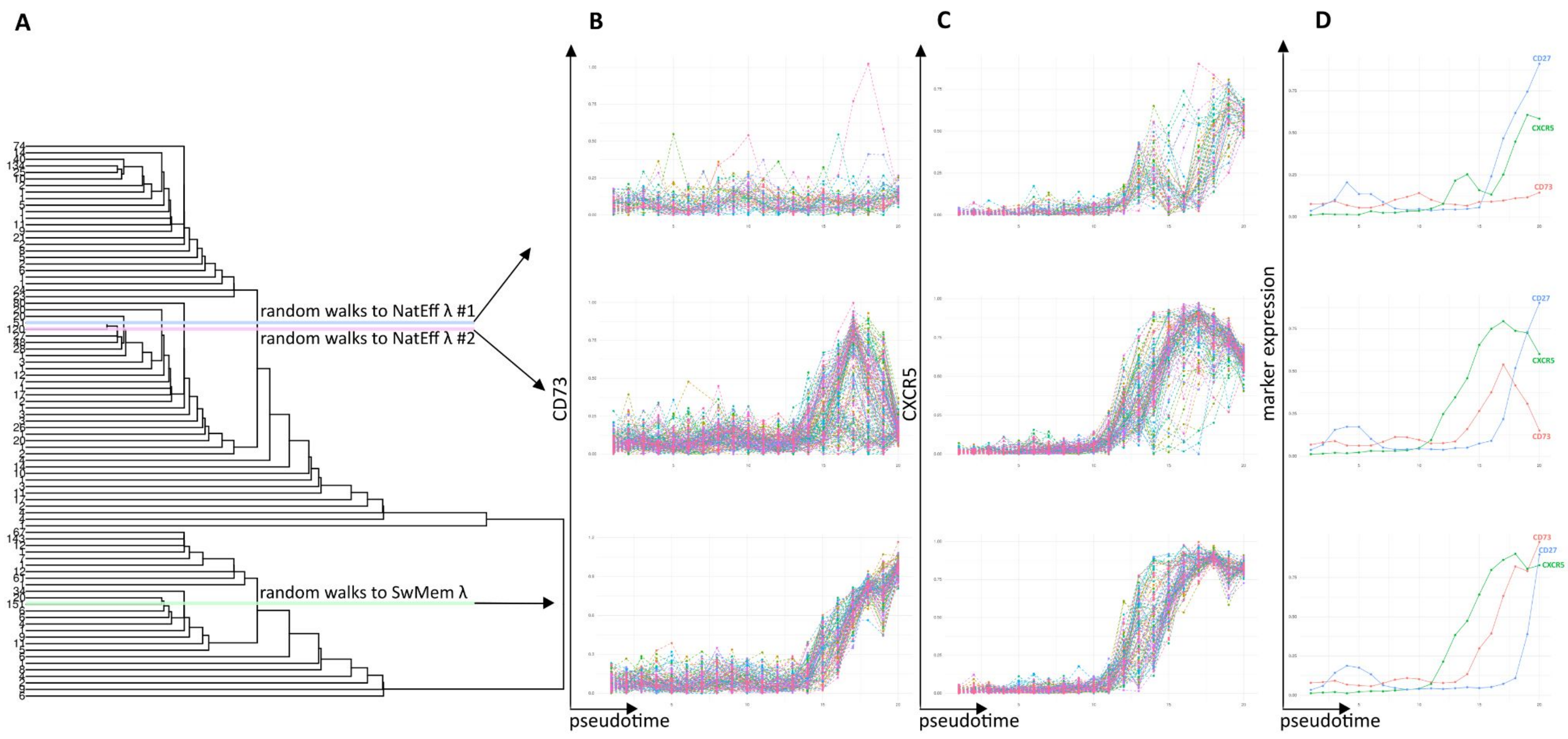


**Supplementary Figure 12.** Differential expression of the markers CD73 and CXCR5 in trajectories leading to Natural Effector and Switched memory cells. (A) Dendrogram with selection of random walks forming the trajectory to Natural Effector λ #1(blue), #2 (pink) and Switched memory λ (green). The trajectory to Natural Effector λ #1 shows no upregulation of CD73 (B, top) and transient expression of CXCR5 (C, top). The trajectory to Natural Effector λ #2 (B, mid) shows heterogeneous expression of CD73 and upregulation of CXCR5 (C, mid). The trajectory to Switched memory λ shows clear upregulation of both CD73 (B, bottom) and CXCR5 (C, bottom). (D) Lineplots for the different trajectories with dynamics of expression of CD73, CXCR5 and CD27.

**Supplementary table 1**.Mass cytometry panel of markers used for staining of the samples. The marker in brackets (iIgM) was excluded from the manual analysis due to poor performance.

| marker | clone | tag | µl per 100 µl cell suspension | manufacturer |
|---|---|---|---|---|
| **BARCODES** | | | | |
| CD45 | HI30 | Y89 | 1 | Fluidigm |
| CD45 | MEM-28 | 110Cd | 2 | Exbio |
| CD45 | MEM-28 | 113In | 1 | Exbio |
| HLA-I | W6/32 | 116Cd | 2 | Bxcell |
| HLA-I | W6/32 | 175Lu | 1 | Bxcell |
| **SURFACE MARKERS** | | | | |
| CD38 | HIT2 | 141Pr | 2 | Exbio |
| sIgM | MHM-88 | 144Nd | 1 | Biolegend |
| CD24 | ML5 | 145Nd | 2 | Biolegend |
| IgD | IA6-2 | 146Nd | 1 | Fluidigm |
| CD20 | 2H7 | 147Sm | 1 | Fluidigm |
| CD34 | 581 | 148Nd | 2 | Fluidigm |
| CD127 | A019D5 | 149Sm | 2 | Fluidigm |
| Ig light chain lambda | MHL-38 | 151Eu | 1 | Fluidigm |
| CD135 | BV10A4 | 156Gd | 4 | Exbio |
| CD10 | HI10a | 158Gd | 1 | Fluidigm |
| CD22 | HIB22 | 159Tb | 2 | Fluidigm |
| Ig light chain kappa | MHK-49 | 160Gd | 1 | Fluidigm |
| CD9 | MEM-61 | 161Dy | 2 | Exbio |
| CD25 | MEM-181 | 162Dy | 4 | Exbio |
| CD44 | MEM-85 | 163Dy | 2 | Exbio |
| CD27 | L128 | 167Er | 1 | Fluidigm |
| CD19 | HIB19 | 169Tm | 2 | Fluidigm |
| CXCR5 | REA103 | 171Yb | 2 | Miltenyi |
| CXCR4 | REA649 | 173Yb | 2 | Miltenyi |
| HLA-DR | L243 | 174Yb | 1 | Fluidigm |
| CD73 | AD2 | 176Yb | 2 | Exbio |
| **INTRACELLULAR MARKERS** | | | | |
| Caspase 3 (Cleaved) | D3E9 | 142Nd | 2 | Fluidigm |
| cPARP | F21-852 | 143Nd | 1 | Fluidigm |
| PAX-5 | 1H9 | 150Nd | 2 | Biolegend |
| Caspase 7 (Cleaved) | D6H1 | 152Sm | 1 | Fluidigm |
| BCL-2 | Bcl-2/100 | 153Eu | 2 | Exbio |
| Tdt | E17-1519 | 164Dy | 1 | Fluidigm |
| Biotin | 1D4-C5 | 165Ho | 1 | Fluidigm |
| Ki-67 | B56 | 168Er | 1 | Fluidigm |
| CD79a | HM57 | 170Er | 2 | Exbio |
| (iIgM | MHM-88 | 172Yb | 1 | Fluidigm) |
| **LINEAGE NEGATIVE MARKERS** | | | | |
| CD3 | UCHT1 | biotin | 2 | Exbio |
| CD16 | 3G8 | biotin | 2 | Biolegend |
| CD33 | HIM3-4 | biotin | 2 | Exbio |
| CD66b | G10F5 | biotin | 2 | Biolegend |
| **DNA INTERCALATOR** | | | | |
| | | 191Ir/193Ir | 1 | Fluidigm |
